# Ascorbic Acid (Vitamin C) Attenuates Dichlorvos-Induced Neurobehavioral and Oxido-Inflammatory Stress in Discrete Brain Regions

**DOI:** 10.64898/2026.10.22.760251

**Authors:** Doris Kasarachi Ogbuokiri, Joshua Izuchukwu Abugu, Olajumoke Blessing Oladosu, Ibrahim Benedicta Unekwuojo, Nkechiyerem Precious Agufusi, Eze Victor Kenechukwu

## Abstract

**Background:** Dichlorvos is a widely used organophosphate pesticide associated with neurotoxicity through acetylcholinesterase inhibition, oxidative stress, and neuroinflammation. Chronic exposure may cause neuronal dysfunction and structural brain alterations. Ascorbic acid (vitamin C), a potent antioxidant, may provide neuroprotection against oxidative damage. This study evaluated the protective effects of ascorbic acid against dichlorvos-induced neurobehavioral and oxido-inflammatory alterations in adult male Wistar rats.

**Methods:** Thirty-six rats were randomly assigned into six groups (n = 6): control, dichlorvos (20 mg/kg), dichlorvos plus low-dose ascorbic acid (50 mg/kg), low-dose ascorbic acid, dichlorvos plus high-dose ascorbic acid (150 mg/kg), and high-dose ascorbic acid groups. Treatments were administered orally for six weeks. Neurobehavioral performance was assessed using the T-maze and hanging-wire tests. Oxidative stress biomarkers, including malondialdehyde, superoxide dismutase, and catalase, were evaluated in the hippocampus and amygdala. Histological examinations of the hippocampus, cerebellum, and prefrontal cortex were also performed. Data were analyzed using one-way ANOVA with Tukey’s post hoc test ().

**Results:** No significant difference in brain weight was observed across groups. Neurobehavioral assessments showed no statistically significant differences between pre- and post-treatment performances, although dichlorvos exposure produced a slight decline, while ascorbic acid-treated groups showed dose-dependent improvement. Dichlorvos significantly increased malondialdehyde levels and decreased superoxide dismutase and catalase activities compared with control (p < 0.05). Co-administration of ascorbic acid significantly reduced lipid peroxidation and improved antioxidant enzyme activities relative to the dichlorvos group (p < 0.05). Histologically, the hippocampus remained largely preserved, while mild neuronal degeneration was observed in the prefrontal cortex of dichlorvos-treated rats. The cerebellum showed mostly normal architecture, with mild molecular layer degeneration in the high-dose dichlorvos plus ascorbic acid group.

**Conclusions:** Ascorbic acid attenuated dichlorvos-induced oxidative stress and reduced associated histopathological alterations, with limited but observable improvements in neurobehavioral outcomes. These findings suggest its potential protective role against organophosphate-induced neurotoxicity.

## INTRODUCTION

Organophosphate pesticides are widely used in households and agriculture for pest control. Environmental exposure to these compounds has emerged as a global health threat, particularly due to their chronic neurotoxicity. Among these chemicals is *dichlorvos* (2,2-dichlorovinyl dimethyl phosphate), popularly known as sniper, commonly used in agriculture and pest control (EPA, 2022). Even though it’s highly efficient in the management of pests and insects, studies have reported its significant effect on mammals by a systemic inhibition of acetylcholinesterase, the enzyme that facilitates the hydrolysis of acetylcholine in the nervous system. The inhibition of acetylcholine triggers the excessive stimulation of cholinergic receptors, which disrupts neuronal signaling, which can contribute to seizures, respiratory failure, and long-term neurological impairments (Mansouri et al., 2021; Bajgar et al., 2020). Dichlorvos triggers oxidative stress by generating reactive oxygen species (ROS), which alters cellular redox homeostasis, thereby leading to lipid peroxidation in cerebellar neurons that leads to cellular dysfunction and apoptosis (Bharathi *et al.,* 2022. The metabolic transformation of this compound yields free radicals that can damage lipids, proteins, and Deoxyribonucleic Acid (DNA) (Zhang *et al.,* 2020). Following prolonged exposure, it can lead to ataxia, tremor, headache, dizziness, and, in extreme cases, longterm cognitive impairment Due to the brain’s lipid-rich membrane and high metabolic demand, the brain is easily vulnerable to neurotoxicity. For instance, dichlorvos has been shown to affect the cerebellum by impairing neuronal activity, thereby undermining its regulatory role in movement and posture. It also impairs the cognitive role of the prefrontal cortex by triggering structural alterations like neuronal shrinkage, loss of Nissl substance, vacuolations, and astrocyte activation (Ogunlade *et al.,* 2017; Kaur *et al.,* 2022). Furthermore, in the hippocampus, dichlorvos exposure is associated with shrunken pyknotic nuclei, significant reduction in the number of viable neurons, and suppression of neurogenesis (Owoeye *et al.,* 2014; Imam *et al.,* 2018). These neuronal effects provide the basis for cognitive deficits and behavioral alterations observed in both humans and experimental models following prolonged organophosphate exposure.

However, exogenous supplementation with ascorbic acid has been discovered to mitigate the neural damage caused by organophosphates by enhancing antioxidant enzyme activity, reducing lipid peroxidation, and improving histological integrity of neural tissues (Njoku *et al.,* 2021; Olajide *et al.,* 2020; Okon *et al.,* 2019). Ascorbic acid generates free radicals, thereby activating other antioxidants like vitamin E, and maintaining the redox balance of neuronal tissues (May et al., 2013; Naidu, 2003). Ascorbic acid is present abundantly in the central nervous system, where it supports neurotransmitter synthesis, protects against oxidative injury, provides immune support, and promotes collagen synthesis (Patel et al., 2018; Carr and Vissers, 2020). Furthermore, ascorbic acid has been revealed to have antioxidant properties that help protect against neuroinflammation and support mitochondrial function (Parab et al., 2024; Cortes-Rodríguez et al., 2023). Although dichlorvos has been associated with cognitive deficit and behavioral impairment, there remains a lack of clarity regarding the extent to which ascorbic acid preserves cortical integrity under exposure to organophosphates. The neuroprotective mechanisms of ascorbic acid, particularly in pesticide-induced neural degeneration, have not been clearly delineated through controlled animal studies. This study aims to investigate the neuroprotective potential of ascorbic acid against dichlorvos-induced neurotoxicity in the brain of adult male Wistar Rats, owing to ascorbic acid’s antioxidant and neuroprotective properties. Ascorbic acid has attracted interest as a potential therapeutic strategy, especially in settings with limited infrastructure where conventional treatments may not be readily available.

## METHODS

### Ethical Approval

Ethical approval was obtained from the ethical committee, Faculty of Basic Medical Sciences, College of Health Sciences, Nnamdi Azikiwe University. NAU/CHS/NC/FBMS/933, This research was carried out at the Department of Anatomy, Faculty of Basic Medical Sciences, College of Health Sciences, Nnamdi Azikiwe University, Nnewi, Nigeria.

### Experimental Animals and Procurement of Dichlorvos and Ascorbic Acid

A total of 36 adult male Wistar rats were obtained for this experiment. The animals were procured from a research enterprise at the University of Ibadan, Oyo State, and housed in the animal facility of the Faculty of Basic Medical Sciences.

The rats were kept in four cages and divided into 6 groups (n = 6); (Control group, Dichlorvos group, Dichlorvos + Low-dose of ascorbic acid group, Low-dose of ascorbic acid group, Dichlorvos + High-dose of ascorbic acid group, and High-dose of ascorbic acid group) The animals were fed with Vital top feed finisher and water for a period of two weeks to enable them acclimatize to their new environment before the commencement of the experiment proper. During the period of acclimatization, water and feed (*ad libitum*) were provided to experimental animals The animal care handling was conducted in compliance with the National Regulation for Animal Research.

Dichlorvos was purchased from Roban stores located at Nnewi-Nnobi Rd, Nnewi 434116, Anambra. The ascorbic acid tablets were purchased from Assumption pharmacy located at 2WQ2+P33 John Cross, Old Oba-Nnewi Rd, Nnewi 435101, Anambra.

### Experimental Procedure

i. **Stock solution for dichlorvos:** A volume of 2 mL of undiluted dichlorvos solution was measured and transferred into a clean storage container. Subsequently, 98 mL of distilled water was added to the same container to obtain a 100 mL stock solution. The mixture was thoroughly homogenized to ensure uniformity. From this stock solution, 0.1 mL was measured and administered.
ii. **Stock solution for ascorbic acid:** A total of 1000 mg of ascorbic acid (equivalent to 10 tablets) was ground into a fine powder on a clean metal plate using a spoon. The powdered sample was then transferred into a clean, dry storage container. Subsequently, 10 mL of distilled water was measured and added to the container, and the mixture was thoroughly homogenized to obtain a uniform solution. From this prepared solution, 0.1 mL was administered for the low-dose group, while 0.4 mL was administered for the high-dose group.

Ascorbic acid was administered orally 30 minutes after dichlorvos had been administered.

### Experimental Design

The experiment lasted for a period of 8 weeks; two (2) weeks for acclimatization and six (6) weeks for administration of dichlorvos and ascorbic acid, and was done using the oral gavage method. Animals were weighed and grouped as follows;

I. Group A (Control group): Rats received normal saline, rats’ chow, and water *ad libitum* for 6 weeks.
II. Group B (Dichlorvos-treated group): Rats were orally administered with 20mg/kg of dichlorvos once daily,
III. Group C (Dichlorvos + Low-dose of Ascorbic acid treated group): Rats were orally administered 20mg/kg of dichlorvos once daily, 50 mg/kg of ascorbic acid once daily,
IV. Group D (Low-dose of Ascorbic acid-treated group): Rats were orally administered 50 mg/kg of ascorbic acid once daily.
V. Group E (Dichlorvos + High-dose of Ascorbic acid treated group): Rats were orally administered 20mg/kg of dichlorvos once daily, 150 mg/kg of ascorbic acid once daily,
VI. Group F (High-dose of Ascorbic acid treated group): Rats were orally administered 150 mg/kg body weight of ascorbic acid once daily,

### T-Maze Test

Neurobehavioral tests are experimental techniques designed to evaluate the relationship between brain function and behavior in animal models or humans (Van der Staay*et al.,* 2009). The T-maze is a widely used behavioral paradigm in neuroscience research to evaluate spatial learning, memory, and decision-making in rodents. It is shaped like the letter ‘T’, consisting of a start arm and two goal arms, which provide the animal with a choice (Deacon and Rawlins 2006). This Tmaze test allows animals to begin at the base and select between two goal arms positioned at the top. When subjected to two consecutive trials, rodents often select the arm not previously visited, an outcome referred to as spontaneous alternation, which reflects short-term memory (Deacon and Rawlins 2006). Both spontaneous and rewarded alternation are highly sensitive indicators of hippocampal dysfunction, although other brain regions also contribute, while each trial generally lasts less than two minutes (Deacon and Rawlins 2006).

### Steps in the T-maze test

#### A. Apparatus Preparation

The T-maze is first prepared by setting it up either in an elevated or enclosed form, consisting of a single start arm (the stem) and two goal arms positioned at right angles to form the shape of the letter “T”. To eliminate any potential olfactory cues that might influence the animal’s behaviour, the maze is cleaned between trials with 70% ethanol or another suitable odour-neutralizing agent (Deacon and Rawlins 2006).

#### B. Testing Phase

During the testing phase, each rodent is placed at the start arm, facing the junction of the maze, and allowed to choose between the two goal arms. In alternation tests, two consecutive trials are conducted in quick succession, with successful performance indicated by the animal choosing the opposite arm on the second trial. Each trial typically lasts less than two minutes to maintain consistency and reduce fatigue (Deacon and Rawlins 2006).

#### C. Recording Behavioural Data

Behavioural data are then recorded for later analysis. Key variables include the arm chosen (left or right), the number of correct alternations, the latency to enter an arm, and the number of errors, such as repeatedly selecting the same arm when an alternation is expected. These observations are compared across experimental groups or conditions to evaluate differences in spatial working memory and decision-making (Deacon and Rawlins 2006).

The T-maze is widely regarded as a sensitive tool that examines the hippocampus, a brain region that plays a central role in spatial learning, memory formation, and decision-making, particularly the hippocampal-dependent spatial learning and memory (Deacon and Rawlins 2006).

The T-maze test was performed pre- and post-administration.

**Fig 3.7.**
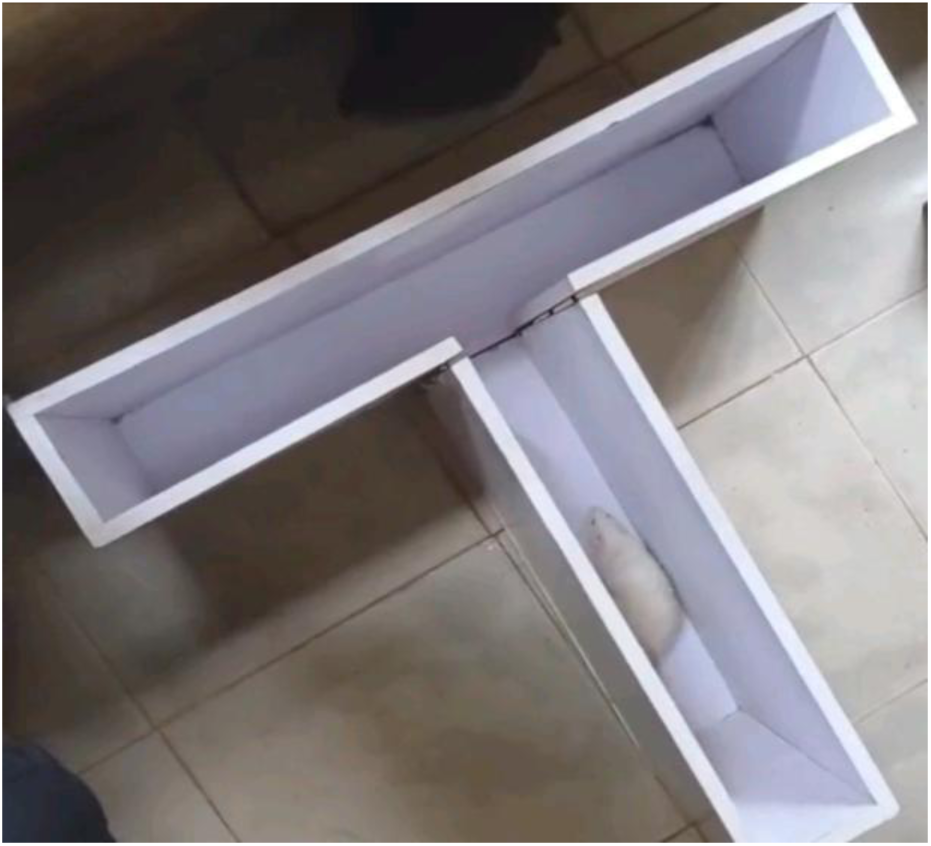
image of the rat at the start point.

### Hanging wire test)

The hanging-wire (also called wire-hang or grid-hang) test is a simple, reliable neurobehavioral assay used to assess forelimb grip strength, endurance, and motor coordination in mice and rat(Shi *et al.,* 2021; Zhang *et al.,* 2023). It is commonly employed to detect deficits resulting from neuromuscular disease, cerebellar dysfunction, or pharmacological interventions that alter motor systems (Jansen-West *et al.,* 2022; Ueno *et al.,* 2022). The test measures an animal’s ability to cling to a suspended wire or inverted grid and is highly sensitive to impairments in balance, coordination, and muscle endurance that frequently accompany cerebellar pathology (Xiao *et al.,* 2020; Zhang *et al.,* 2023).

The hanging-wire test was conducted to assess forelimb grip endurance and motor coordination of the adult male wistar rat. A single stainless-steel wire (2 mm diameter, length ∼55 cm) was mounted horizontally between two upright supports and positioned at a height of ∼35 cm above a soft foam padding to prevent injury in the event of a fall. Prior to each session the apparatus was wiped with 70% ethanol using cotton wool to remove olfactory cues. Each rat was gently held by the tail and lowered until its forepaws grasped the wire; timing commenced immediately upon release. Trials lasted until the rat fell. Each rat underwent three trials with 5-minute inter-trial intervals. The mean latency to fall across the three trials was calculated for each animal and used data analysis.

### Assessment of Brain Superoxide Dismutase (SOD), Catalase (CAT), Glutathione S-Transferase (GST) Activities, and MDA Levels

#### Measurement of Oxidative Stress

The present study focused on several key parameters, including malondialdehyde (MDA), GSH, and SOD. These parameters were measured using specific kits from ZellBio (Germany), and quantitation of OS in biological samples was conducted according to the manufacturer’s protocols. The MDA assay (CAT No. ZB-MDA-96A) was used to assess lipid peroxidation, indicating oxidative damage. The GSH assay (CAT No. ZB-GSH-96A) measures the levels of this critical antioxidant, Finally, the SOD assay (CAT No. ZB-SOD-96A) evaluates the activity of this enzyme that protects against OS. Following the manufacturer’s instructions for each assay ensured accurate and reproducible results, contributing to the understanding of the biochemical changes associated with ethanol toxicities and the potential therapeutic effects of the treatments administered

### Sample preparation

**After the completion of the experimental procedures (after 3 weeks), the animals were humanely sacrificed in accordance with ethical guidelines to minimize suffering. This process typically involved administering with** chloral hydrate (300mg/kg, *i.p.*) and perfused with 0.1M phosphate-buffered saline**, ensuring that the animals were unconscious and pain-free before euthanasia. Following sacrifice, Amygdala and Hippocampal tissues were collected for further analysis. The** hippocampus and amygdala **tissues were carefully excised and stored in sterile containers,** The isolated hippocampus and amygdala were weighed and homogenized in 0.03 M sodium phosphate buffer, *p*H-7.4 with an Ultra-Turrax T25 (USA) homogenizer at a speed of 9500 rpm. The hippocampus and prefrontal cortex homogenate was kept at -20°C.

### Statistical Analysis

We used IBM® Statistical Package for the Social Sciences Statistics software (ver. 20) to analyze data, and the results are expressed as mean ± standard deviation. A one-way analysis of variance was conducted followed by Tukey’s post hoc test to calculate the differences between groups.

A P < 0.05 was considered to indicate statistical significance.

## RESULTS

### Effect of Ascorbic Acid and Dichlorvos on the Rats’ Brain Weight

The brain weight of Wistar rats was obtained and measured 24 hours after the final treatment.

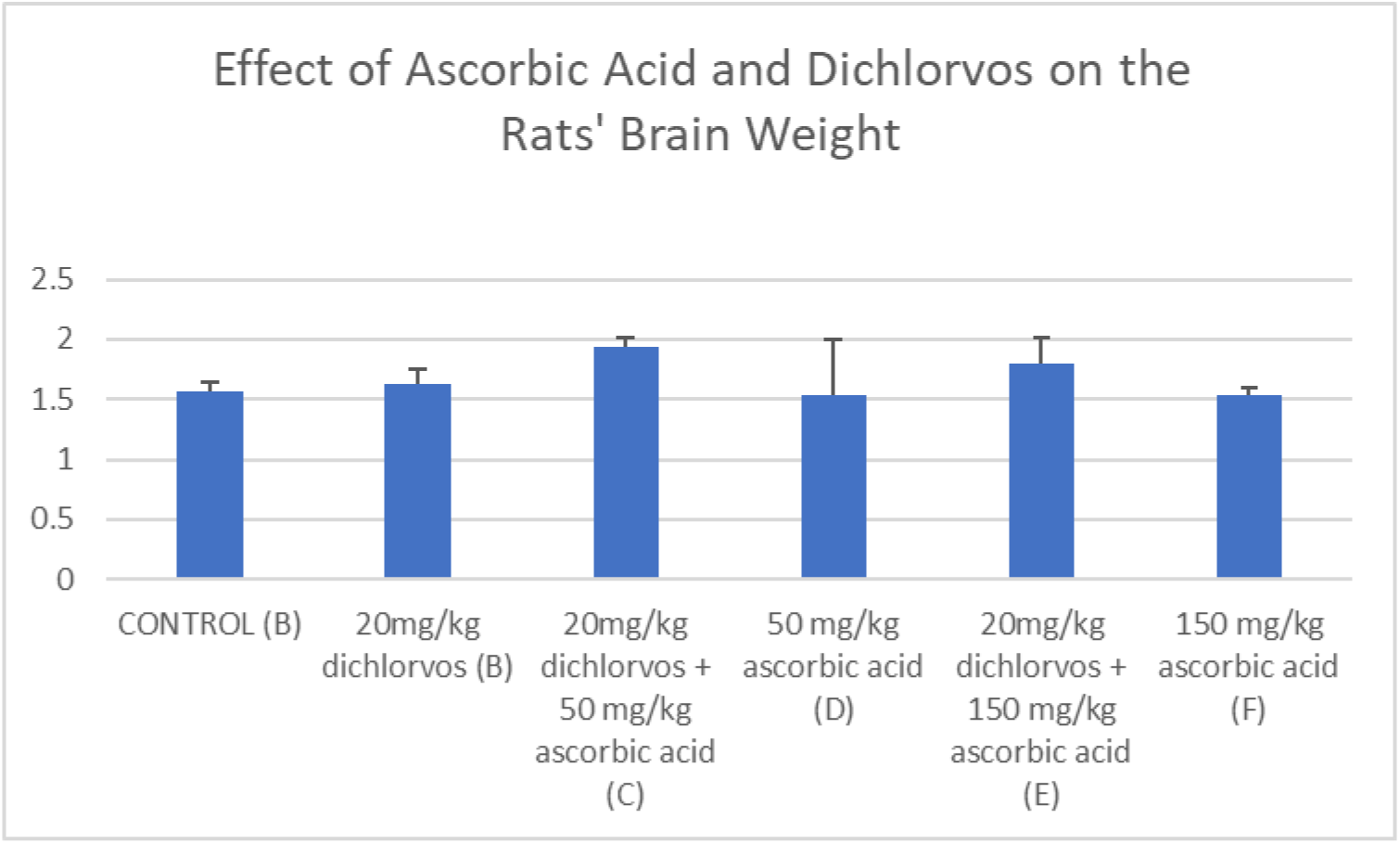
Data were analyzed using One-way ANOVA, followed by Post HOC Tukey’s multiple comparison, and data were considered significant at P<0.05

The result shows no statistically significant difference in the brain weight **T-maze test analysis**

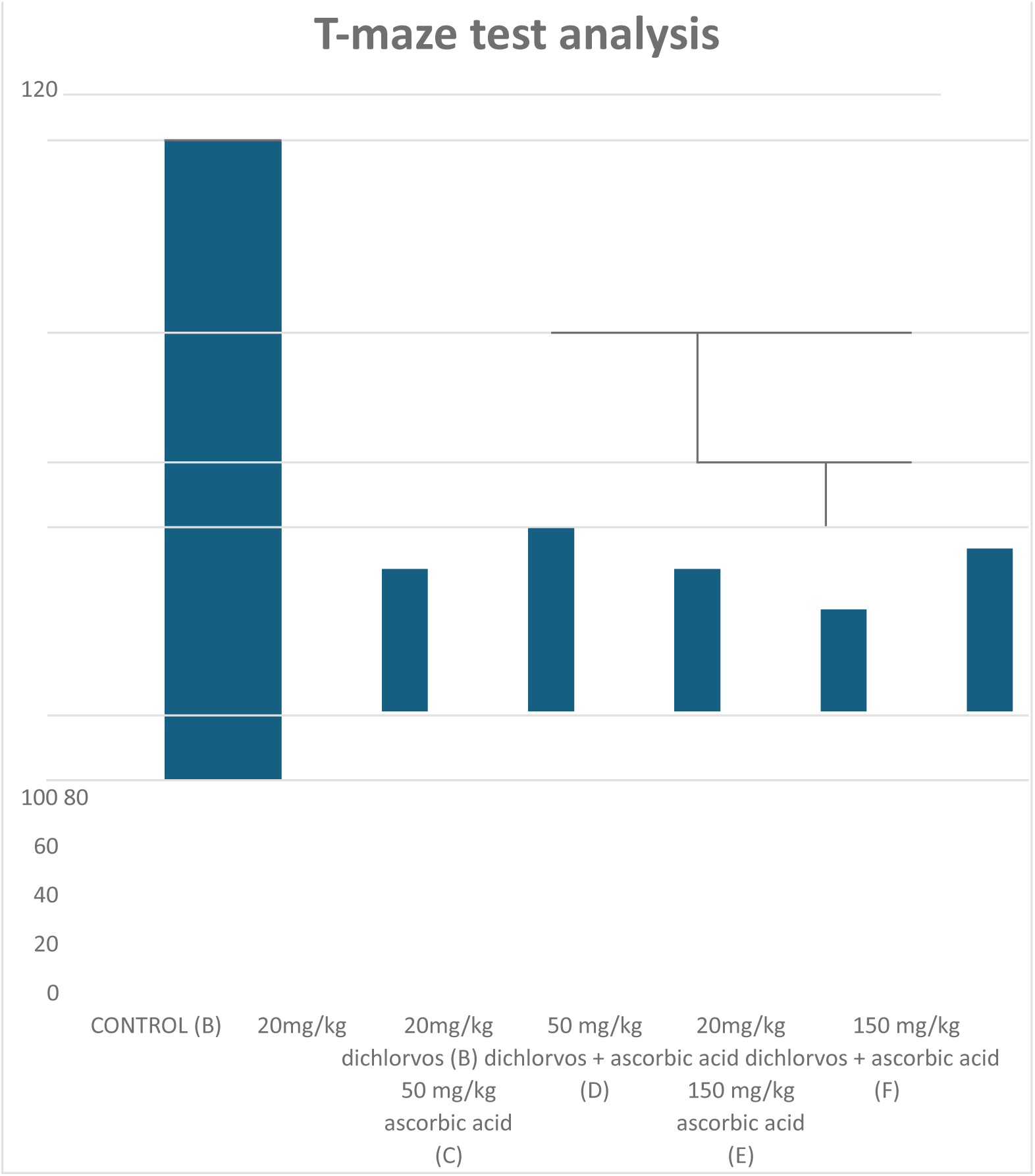
Data were analyzed using One-way ANOVA, followed by Post HOC Tukey’s multiple comparison, and data were considered significant at P<0.05

Result shows that there was no significant difference in the T-Maze test, when comparing the initial, test to the final test in the groups exposed to dichlorvos and treated with ascorbic acid, Group B, subjected to dichlorvos only showed non-significant decrease (P<0.05) when comparing the initial test to the final test Group C-F, showed dose dependent differences (P<0.05) when comparing the initial test to the final test.

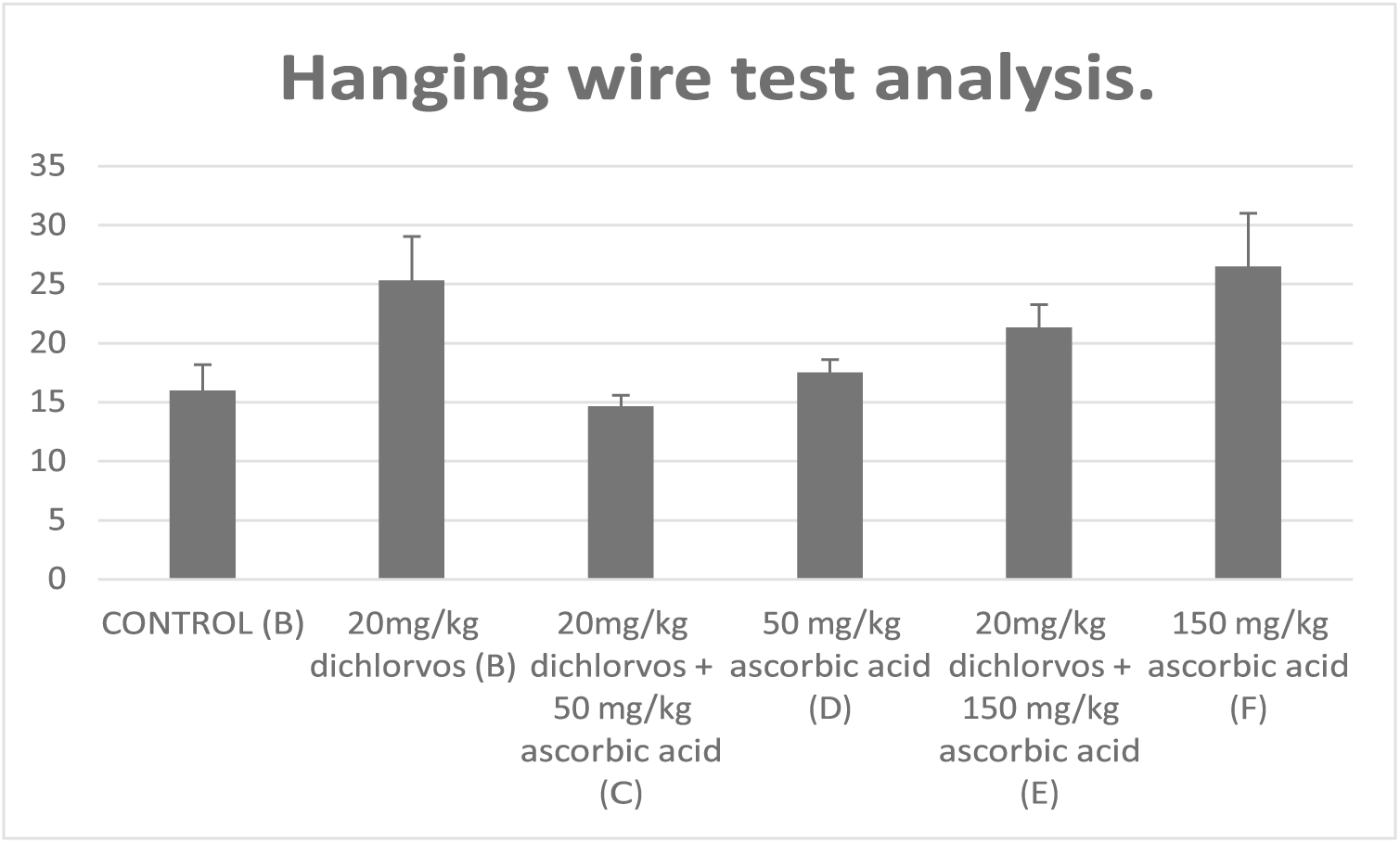
*Data were analyzed using One-way ANOVA, followed by Post HOC Turkey’s multiple comparison, and data was considered significant at P<0.05* (*= significant)

Result from the table above shows that there was no significant difference in the hanging wire test, when comparing the initial test to the final test In the groups exposed to **dichlorvos** and ascorbic acid, Group B, subjected to **dichlorvos** showed non significant decrease (P<0.05) when comparing the initial test to the final test Group, while group C-F, showed dose dependent increase(P<0.05) when comparing the initial test to the final test

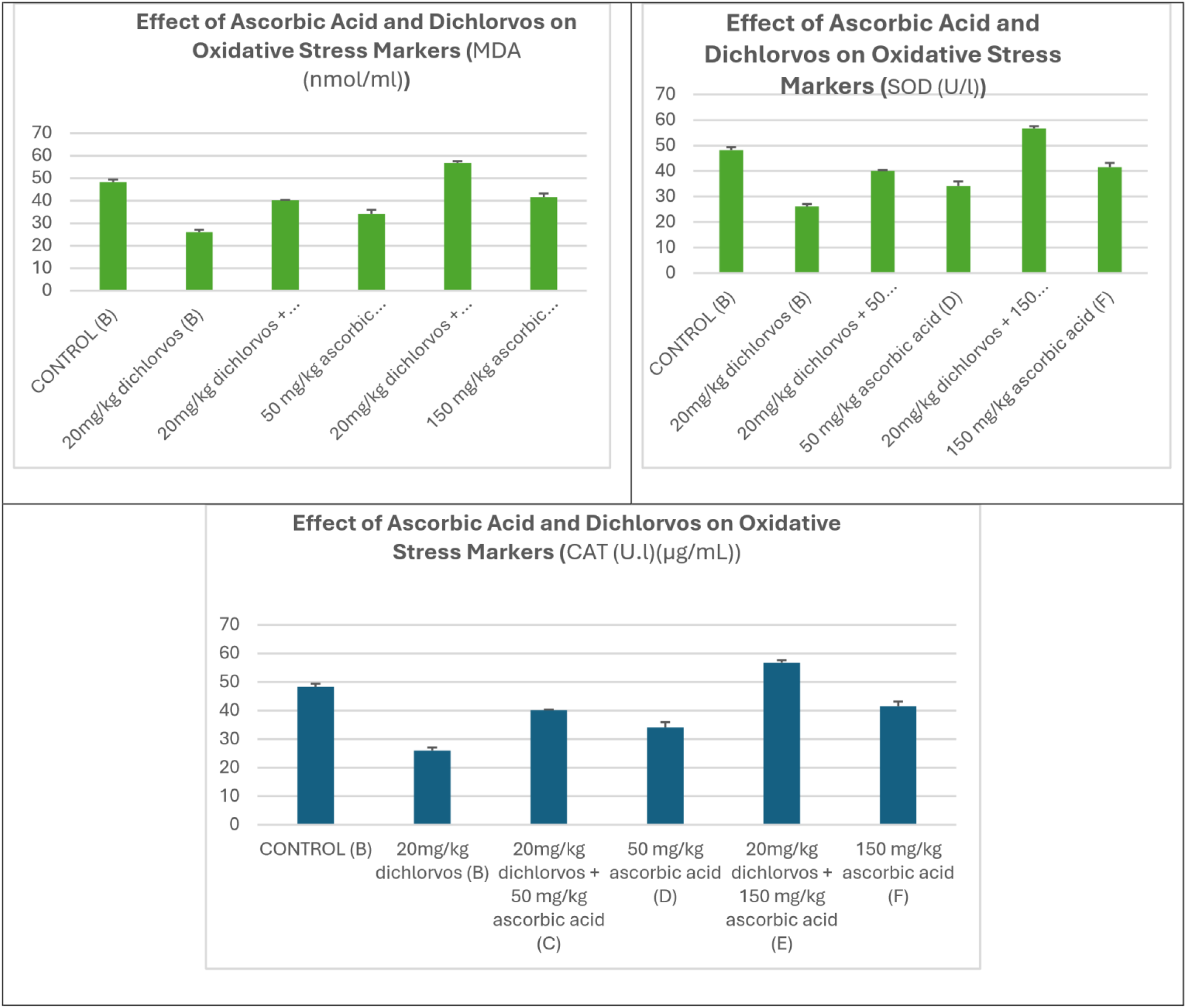
Data were analyzed using One-way ANOVA, followed by Post HOC Tukey’s multiple comparison, and data were considered significant at P<0.05

Results showed a statistically significant increase in the MDA level in group B (dichlorvos) compared to the control (group A), while groups C-F presented a significant difference when compared to group A. The SOD and CAT showed a statistically significant decrease in group B (dichlorvos) compared to control group A, while supplementation of ascorbic acid in groups C-F showed a significant increase when compared to the control.

MDA=Malondialdehyde, SOD= Superoxide dismutase, CAT=Catalase

### Histological Findings

#### hippocampus

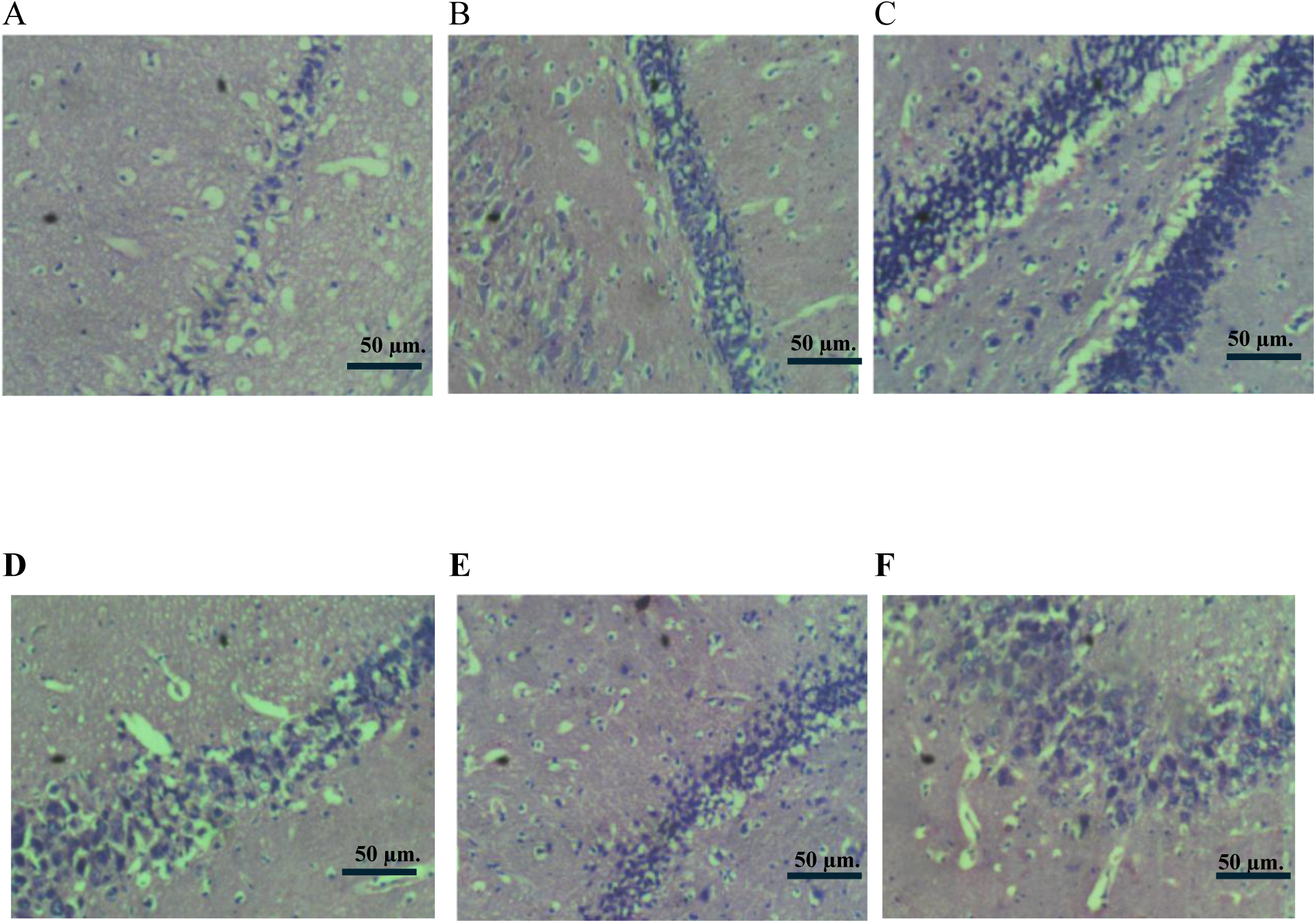

Photomicrographs of the hippocampus consistently show normal cytoarchitecture with densely packed granular neurons of the dentate fascicle and larger histologically intact neurons of the endplate, with no evidence of injury (H and E x400

#### Cerebellum

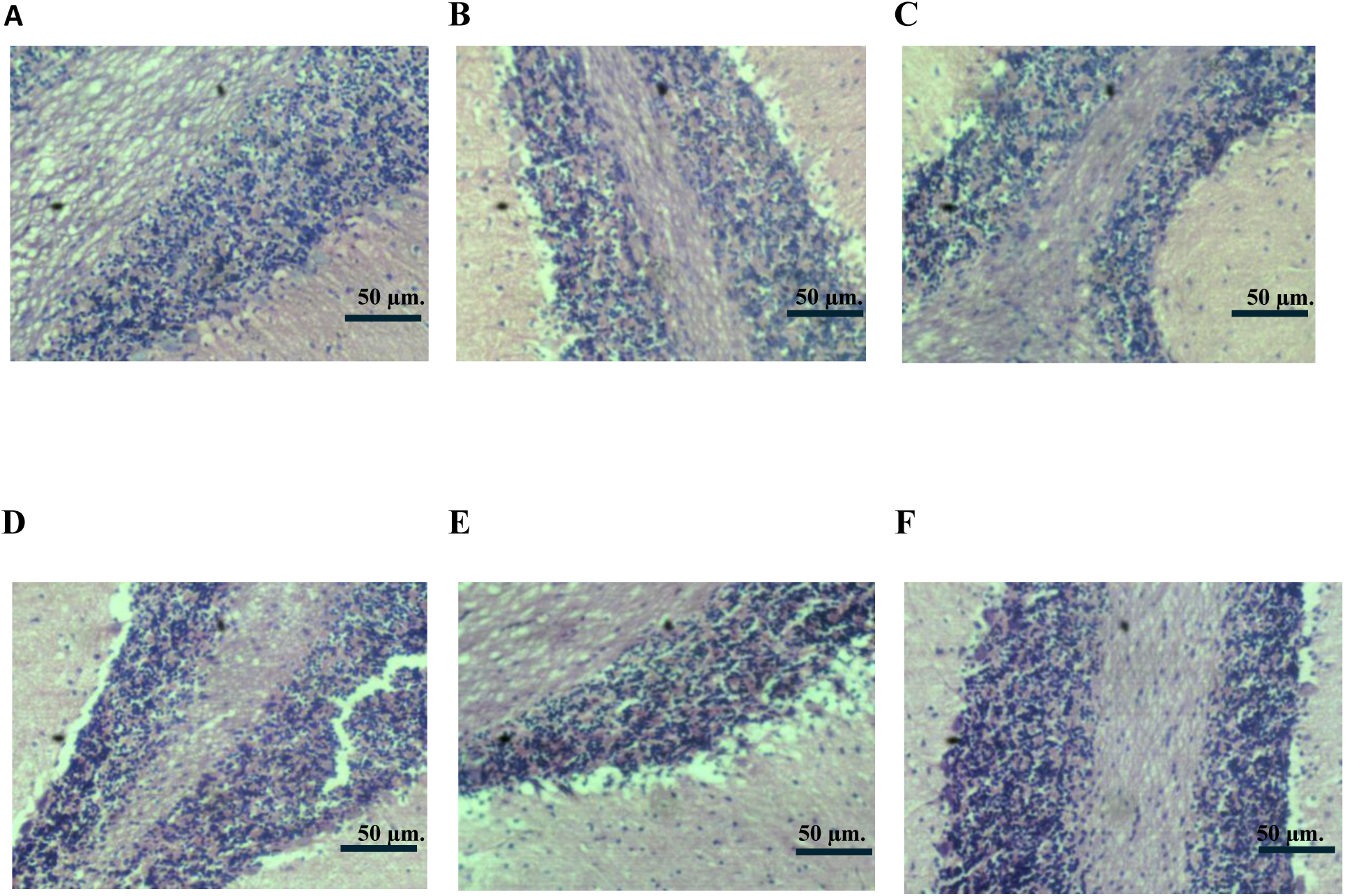

Photomicrographs of brain sections across all groups show the cerebellum with generally normal morphology and intact molecular layer Purkinje fibre layer, and white matter except in the highdose ascorbic acid and dichlorvos group, which exhibits degeneration of the molecular layer despite preserved integrity of other structures (Haematoxylin & Eosin x400).

#### Prefrontal Cortex

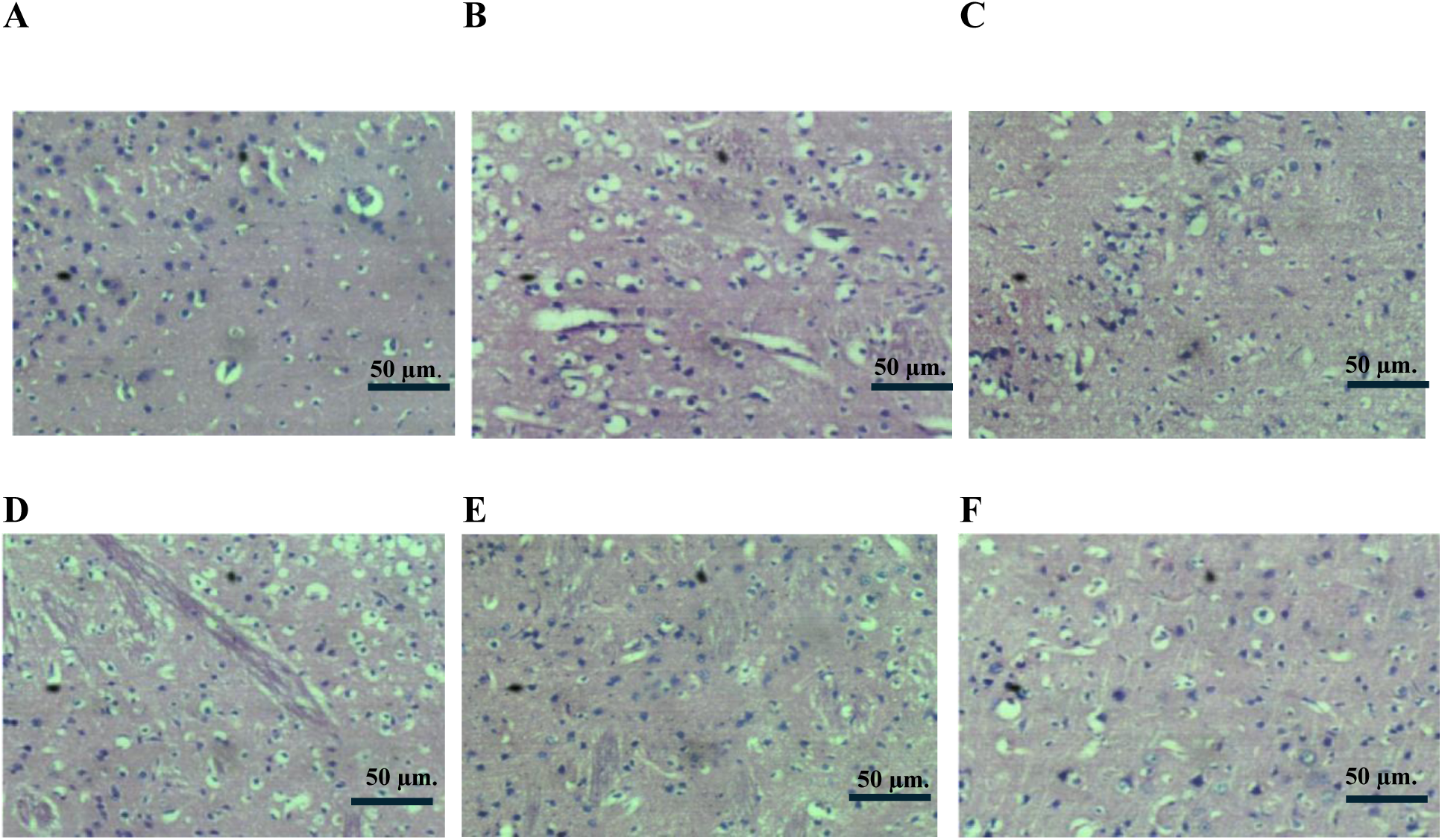

Photomicrographs of the prefrontal cortex across all groups show generally normal histoarchitecture with distinct cortical layers and cellular components, with no inflammatory changes observed in control and high-dose ascorbic acid-treated groups, while mild early neuronal degeneration is noted in the dichlorvos group, mild neuronal and glial degeneration in the low-dose ascorbic acid plus dichlorvos group, and mild connective tissue hypertrophy in the low-dose ascorbic acid group (H and E X400).

## DISCUSSION

Organophosphate pesticides, such as dichlorvos, have been confirmed in several studies to be highly toxic to the mammalian brain. This study further demonstrates that persistent exposure to dichlorvos induces diverse neurotoxicity in the cerebellum, hippocampus, and prefrontal cortex of the adult male Wistar rats. Dichlorvos exerts its toxicity by irreversibly inhibiting acetylcholinesterase (AChE), leading to cholinergic overstimulation and oxidative stress (Mansouri et al., 2021; Bajgar et al., 2020). This study reveals that the administration of ascorbic acid exerts significant neuroprotective effects by enhancing antioxidant enzyme activity, reducing oxidative stress, and preserving normal behavioral responses.

A consistent observation amongst all dichlorvos-induced groups was a reduction in weight compared to the control group, which is indicative of systemic xenobiotic stress. This phenomenon is consistent with previous findings that organophosphate compounds impair metabolic balance and feed intake due to gastrointestinal toxicity and systemic oxidative stress (Yadav et al., 2021; Eze et al., 2020; Smith et al., 2021; Dharmani et al., 2021). However, co administration of ascorbic acid significantly improved body weight gain in a dose dependent manner (Demerdash et al., 2012); Airaodion et al., 2019; Ghorbani-Taherdehi et al.,2020), By mitigating the systemic oxidative stress and improving metabolic efficiency, ascorbic acid likely counteracted the catabolic state induced by dichlorvos, allowing for more normal growth It is important to note that ascorbic acid is a cofactor for numerous metabolic enzymes and is essential for the synthesis of carnitine, which is crucial for fatty acid oxidation and energy production (Lykkesfeldt and Tveden-Nyborg, 2023). The significant weight gain in the ascorbic acid-only groups further supports its role as a vital nutrient promoting overall health and growth. This suggests that ascorbic acid mitigates systemic oxidative stress and restores metabolic efficiency, as revealed in previous studies identifying ascorbic acid as a potent antioxidant capable of reducing pesticide-induced oxidative stress and preventing associated weight gain by neutralizing free radicals and improving hepatic and mitochondrial function (Demerdash et al., 2012). The restoration of weight gain across all brain regions strengthens the evidence that ascorbic acid exerts a systemic protective effect beyond localized neural protection.

While dichlorvos is known to interfere with neuronal metabolism and structural integrity of the brain, potentially leading to atrophy or weight reduction (Assini, Diniz, and Soares, 2017). The findings across the cerebellum, hippocampus, and prefrontal cortex revealed no significant difference in relative brain weight among the groups. Interestingly, there was a slight increase in the dichlorvos-exposed group cotreated with ascorbic acid. A similar pattern of stability in brain weight was reported by Sule (2022), who observed non-significant changes in brain weight following organophosphate exposure but noted histological protection with antioxidant supplementation. The weight of the brain was maintained in the ascorbic acid-treated groups, which suggests the potential neuroprotective effect against the neurotoxicity of dichlorvos, which may be attributed to reduced lipid peroxidation and improved antioxidant enzyme activities. The brain maintenance also indicates that ascorbic acid may not directly influence cerebellar mass within the short experimental duration, although it has the ability to preserve neural tissue integrity under neurotoxic conditions.

There was also a significant increase in malondialdehyde levels in all brain regions, confirming enhanced lipid peroxidation and membrane damage due to dichlorvos exposure. This result mirrors a previous study that revealed that organophosphate exposure increases reactive oxygen species (ROS) production and disrupts cellular antioxidant defense systems (Uchendu et al., 2018), while ascorbic acid maintains antioxidant enzyme activities and reduces lipid peroxidation, validating its efficacy as an effective free-radical scavenger. Study from (Mudgal, 2023) further confirms that ascorbic acid effectively restored redox equilibrium and enhanced endogenous antioxidant defense systems.

Additionally, a substantial decrease in superoxide dismutase (SOD) and catalase (CAT) activities was observed in the dichlorvos-exposed group, indicating a compromised antioxidant defense. In the co-administered group, the restoration of SOD and CAT activities was highly significant; also, CAT activities not only returned to baseline but also increased significantly beyond control levels. This result suggests that ascorbic acid functions not only as a direct scavenger but may also upregulate or preserve the activity of these essential endogenous enzymes.

Also, ascorbic acid has regenerative abilities and has been shown to facilitate the regeneration of other antioxidants such as glutathione, therefore enhancing the functional efficiency of SOD and CAT, including maintaining the synergistic antioxidant network (Akbari et al., 2022). Elevating the dose of ascorbic acid restored the function of these enzymes, which implies a dose-dependent therapeutic effect, underscoring the significance of precise dosing in antioxidant therapy. Markers of antioxidant activity were also restored across the cerebellum, hippocampus, and prefrontal cortex, further validating oxidative stress as the fundamental pathological mechanism and reinforcing the potential of antioxidant therapy as a viable intervention.

Wistar rats subjected to dichlorvos exposure displayed diminished cognitive function in the Tmaze behavioral assessment, as evidenced by reduced percentages of correct responses. The reduction in cognitive function is consistent with the known effects of cholinergic impairment and oxidative stress within the prefrontal cortex (Patel and Jaiswal, 2020). Interestingly, the coadministration of ascorbic acid significantly improved performance, signifying improved memory retention and spatial learning capabilities. The neuroprotective abilities of Ascorbic acid could be attributed to its involvement in neurotransmitter synthesis, the modulation of synaptic activity, and the mitigation of neuronal apoptosis (Harrison and May, 2009). However, in contrast, the hanging wire test results showed no statistically significant differences across all treatment groups. Group exposed to dichlorvos alone exhibited a mild reduction in performance compared to the control, suggesting possible neuromuscular weakness. Although these changes were not statistically different, they displayed consistent directionality across groups. Interestingly, in co-administered group, administration of ascorbic acid, whether at a low dose or high dose, did not produce any significant improvement in motor coordination. In fact, the lower dose group showed a slight, non-significant increase, while the higher dose group demonstrated a mild decrease in motor coordination. This suggests several possibilities, including that oxidative stress is not the sole mechanism of dichlorvos toxicity, but primarily by the inhibition of acetylcholinesterase, leading to cholinergic dysfunction, which ascorbic acid does not directly reverse (Liang et al., 2021). Furthermore, research has shown that organophosphates irreversibly inhibit acetylcholinesterase, causing accumulation of acetylcholine at the neuromuscular junction, resulting in nicotinic receptor stimulation leading to muscle weakness and motor deficits (Angela et al., 2023). Overall, this outcome may reflect that the duration or the dosage of antioxidant administration was insufficient to produce measurable functional recovery, consistent with studies that short-term antioxidant therapy may not immediately translate to behavioral improvement (Afolabi et al., 2021; Sule, 2022).

Histopathology analysis revealed various effects across brain regions. The cerebellum and hippocampus largely maintained their tissue architecture when stained with H&E. In contrast, the prefrontal cortex showed some signs of neuronal damage after exposure to dichlorvos. This difference between biochemical results and histological findings is not unusual in studies of toxins, especially those looking at short-term exposure. It suggests that the dichlorvos-induced oxidative damage at the dose and duration used was primarily at a molecular and synaptic level, preceding gross structural cell death (Uttara et al., 2021). In some instances, high-dose ascorbic acid combined with dichlorvos produced mild molecular layer alterations, suggesting a potential prooxidant interaction under pharmacological concentrations, thereby emphasizing the importance of dose regulation in antioxidant therapy under toxicant exposure (Burgos-Morón et al., 2020; Duarte and Lunec, 2022; Valko et al., 2021). In a recent study, it was revealed that in the presence of transition metals like iron or copper, high concentrations of ascorbic acid may promote redox cycling and ROS formation, which may account for the mild structural alteration and inconsistent behavioral improvements observed in some high-dose groups (Timoshnikov et al., 2020)

A key observation is the dual role of ascorbic acid as an antioxidant and prooxidant depending on the dose. Ascorbic acid demonstrated antioxidant effect at moderate doses, while at higher doses it demonstrated potential pro-oxidant effects which was similarly found in a previous study (Chakrabothy et al., 2014; Chakrabothy et al., 2017).

Techniques like H&E staining might lack the sensitivity to efficiently detect subtle neuronal or synaptic degeneration; it is advisable to utilize more sensitive methods like immunohistochemistry for caspase-3 or synaptophysin in future investigations.

Finally, this study reveals that exposure to dichlorvos induces oxidative, neuronal damage, and behavioral impairment in adult male Wistar rats, but the coadministration of ascorbic acid, particularly at higher doses, substantially mitigates these toxic effects by improving antioxidant enzyme activity, reducing lipid peroxidation, and preserving the histological integrity of the brain. Interestingly, this study underscores the neuroprotective capacity of ascorbic acid against organophosphate-induced neurotoxicity, suggesting its possible therapeutic application in preventing cognitive and structural brain deficits associated with pesticide exposure.

## Conclusion

Overall, ascorbic acid exhibits neuroprotective properties primarily through its antioxidant action, indicating its potential as a therapeutic adjunct in mitigating organophosphate-induced neurotoxicity. Further studies incorporating longer exposure periods, larger sample sizes, and more sensitive behavioral paradigms are recommended to fully elucidate its functional benefits and underlying mechanisms.

## Author Contributions

- **DKO**: Conceptualization, supervision, study design, critical revision of manuscript, project administration. **JIA**: Conceptualization, methodology, investigation, data curation, formal analysis, writing—original draft, visualization. **OBO**: Methodology, investigation, biochemical assays, validation. **IBU**: Investigation, animal handling, behavioral testing, data collection. **NPA**: Investigation, histological analysis, data interpretation. **EVK**: Formal analysis, statistical analysis, manuscript review and editing.

